# Mathematical Modeling Of Systems Biology

**DOI:** 10.1101/2022.08.17.504297

**Authors:** Aaditya Prasad Gupta

## Abstract

A modeling is a mathematical tool, like a microscope, which allows consequences to logically follow from a set of assumptions by which a real world problem can be described by a mathematical formulation. It has become indispensable tools for integrating and interpreting heterogeneous biological data, validating hypothesis and identifying potential diagnostic markers. The modern molecular biology that is characterized by experiments that reveal the behaviours of entire molecular systems is called systems biology. A fundamental step in synthetic biology and systems biology is to derive appropriate mathematical model for the purposes of analysis and design. This manuscript has been engaged in the use of mathematical modeling in the Gene Regulatory System (GRN). Different mathematical models that are inspired in gene regulatory network such as Central dogma, Hill function, Gillespie algorithm, Oscillating gene network and Deterministic vs Stochastic modelings are discussed along with their codes that are programmed in Python using different modules. Here, we underlined that the model should describes the continuous nature of the biochemical processes and reflect the non-linearity. It is also found that the stochastic model is far better than deterministic model to calculate future event exactly with low chance of error.

## 1 Introduction

Mathematical modeling is a process by which a real world problem is described by a mathematical formulation. However, neither all details of single processes will be described nor all aspects concerning the problem will be included. Models are designed to focus on certain aspects of the objects of study. A simple sketch of the mathematical modeling of systems biology is shown in figure 1 which is inspired from the reference^1^. The current modern molecular biology that is characterized by experiments that reveal the behaviours of entire molecular systems is called systems biology^2^.

**Figure 1.**
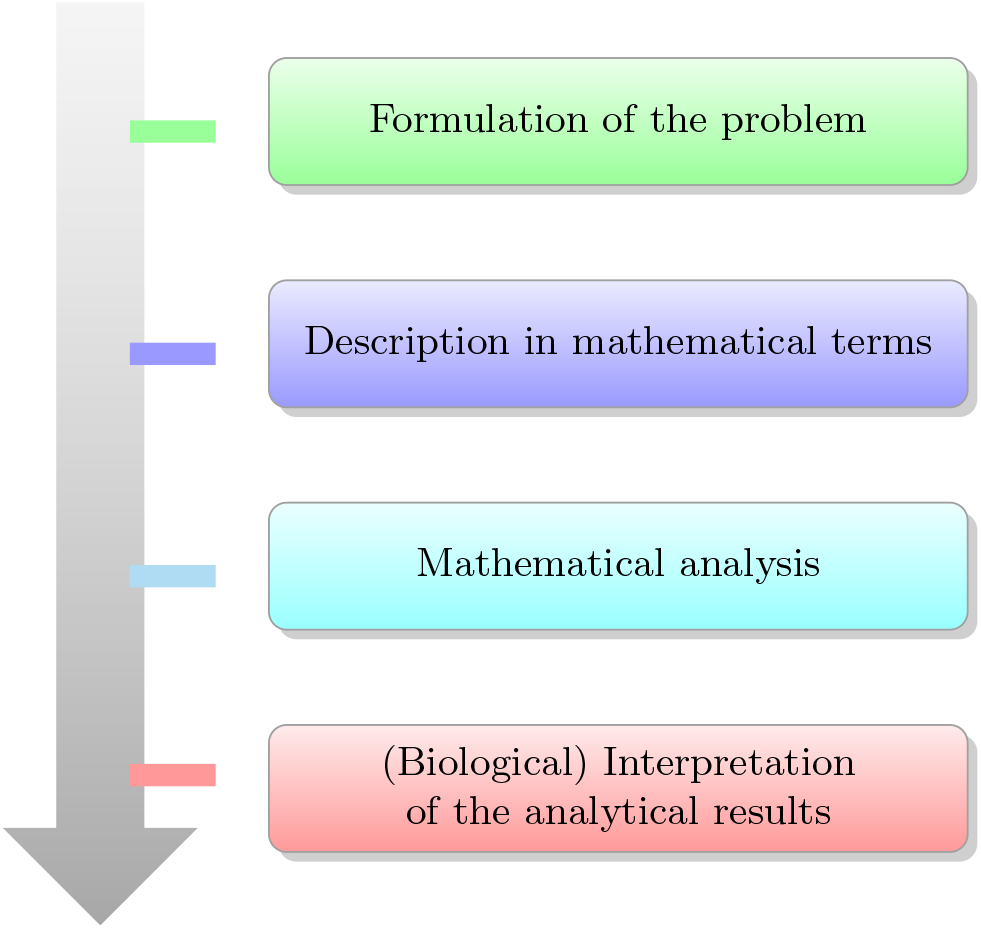
Mathematical formulation for interpreting the analytical biological result

A main problem in systems biology is to find an appropriate mathematical formulation. Then, for further studies of the model, common mathematical tools can be used or new ones are developed. Further, a model should be as simple and detailed as necessary or vice versa. One mathematical formulation may be appropriate for several real-world problem, even from very different domains.

A great challenge of modelling is to bring together the abstract, mathematical formulation and concrete experimental data. The modelling process can be roughly described as shown in figure 2. The figure 2 is inspired from the reference^3^.

**Figure 2.**
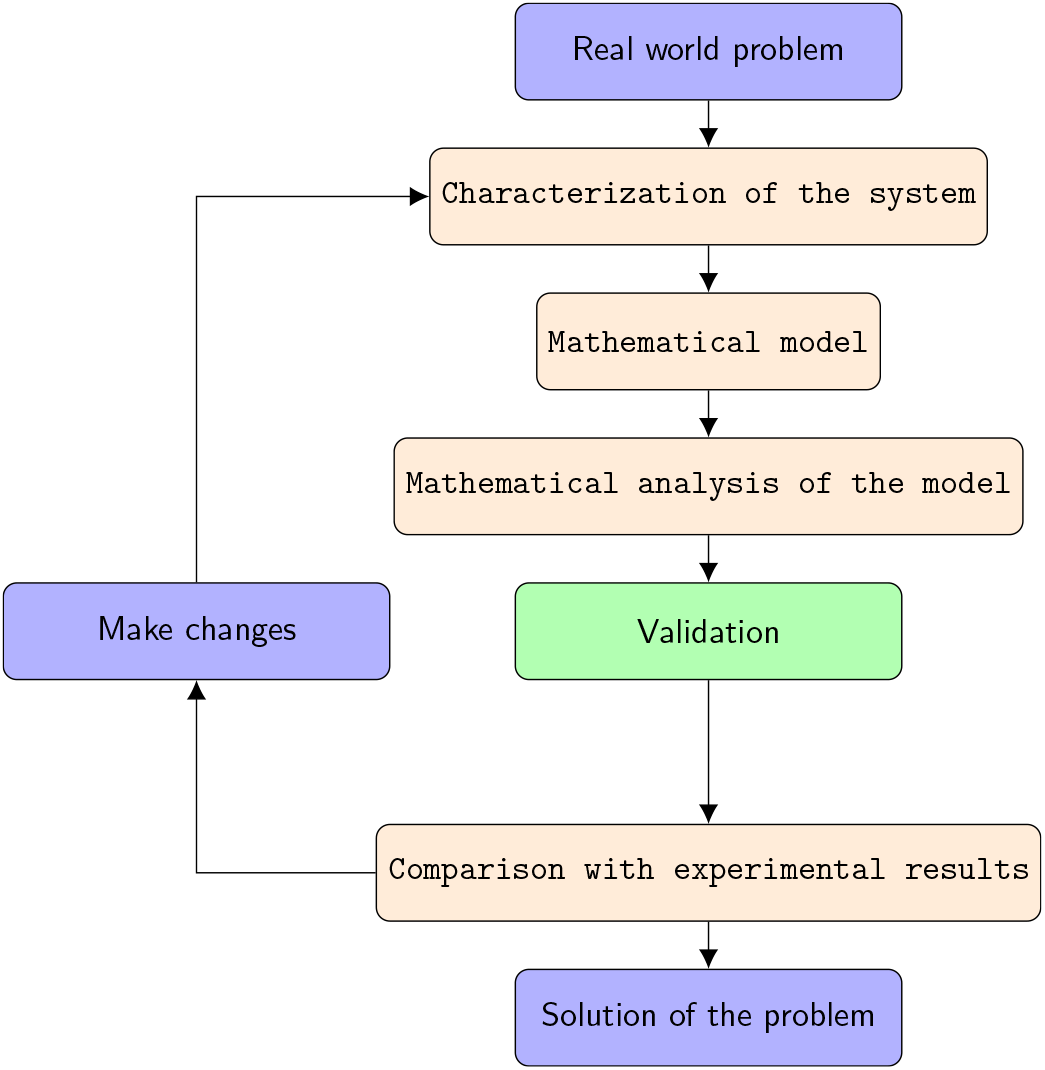
Mathematical model to solve the real world problem

Mathematical modeling has been applied to biological systems for decades, but with respect to gene expression, too few molecular components have been known to build useful, predictive models. There are many different modelling approaches and their number is still increasing. Some of the very famous models in molecular biology are discussed in this manuscript. In section 2 we discuss about the Central dogma. In section 3 Hill function is discussed while Oscillating gene network in section 4, Gillespie algorithm in section 5 and Deterministic vs Stochastic modelings in section 6. Finally, the aspects and scope of mathematical modeling to the above stated models are discussed in section 7. The codes used for the modelling are shown in the appendix along with a brief explanation and a much detailed version on github^1^.

## 2 Central Dogma

Central dogma of molecular biology is an explanation of the flow of genetics information within a biological system. It is often stated as “DNA makes RNA and RNA makes proteins”^4^. The transfer of information from nucleic acid to nucleic acid, or from nucleic acid to protein may be possible but transfer from protein to protein or from protein to nucleic acid is impossible. Information means here precise determination of sequence either of bases in the nucleic acid or of amino acid residue in protein.

The dogma is a framework for understanding the transfer of sequence information between information-carrying bio-polymers in the most common or general case in living organism. There are 3 major classes of such bio-polymers: DNA and RNA (both nucleic acids) and protein. There are 3*3 = 9 conceivable direct transfer of information that can occur between these. The dogma classes these into 3 groups of 3^5^:

- 3 general transfers(believed to occur normally in most cells),
- 3 special transfers(known to occur, but only under specific conditions in case of some viruses or in a laboratory),
- 3 unknown transfers(believed never to occur).

The general transfers describe the normal flow of biological information: DNA can be copied to DNA (DNA replication), DNA information can be copied into mRNA, (transcription), and proteins can be synthesized using the information in mRNA as a template (translation).

The coupled differential equations representing the model of Central dogma are shown in equation 1 & equation 2.

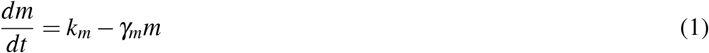

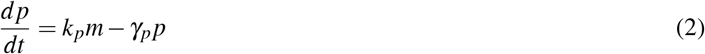

where

- p is protein
- m is mRNA
- *k_m_* is the production rate for the mRNA
- *γ_m_* is the degradation rate for mRNA
- *γ_p_* is the degradation rate of protein
- *k_p_* is the production rate for the proteins

The figures 3 & 4 show the relation between abundance of mRNA and protein over time. Along the y-axis, we have the time axis and along x-axis we have abundance of both mRNA and protein. We observe that M(blue line) denotes mRNA in the above figure 3 & 4 while P(red line) denotes protein. In figure 3 both of them starts at zero and then they increases until they reach their steady state. The steady state for mRNA is 4 and for protein is 16 above. The abundance of protein is more than mRNA because it depends upon the mRNA production (*k_p_*m*) while mRNA production is independent (*k_m_*).

**Figure 3.**
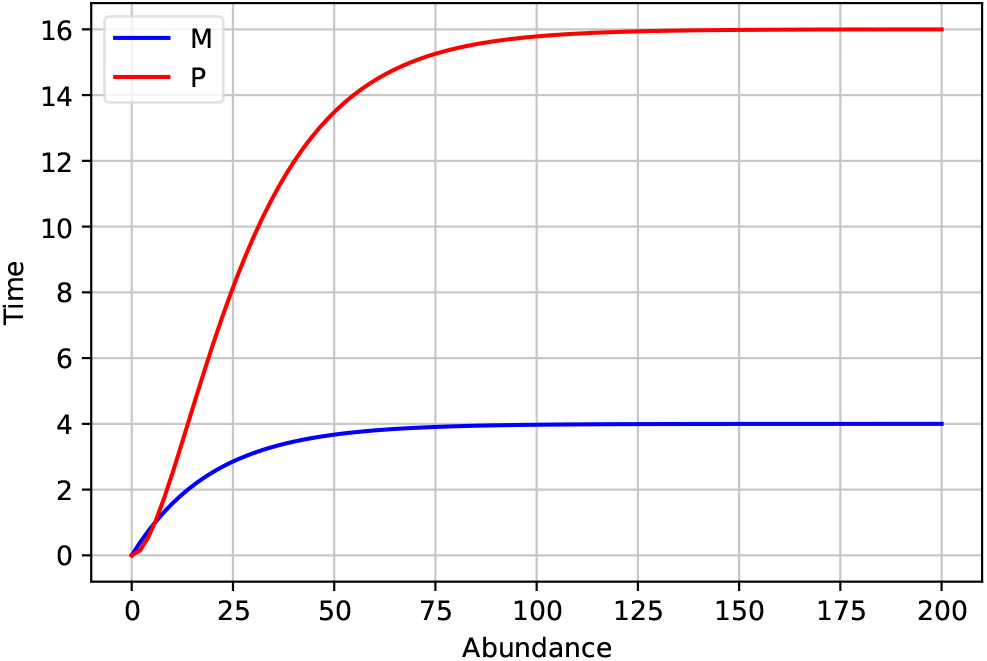
Abundance of mRNA over time with the initial conditions: *k_m_* = 0.2, *γ_m_* = 0.05, *k_p_* = 0.4, *γ_p_* = 0.1

If we change the initial parameters such as *k_m_*, *γ_m_*, *k_p_*, *γ_p_* greater than that are used in figure 3, we came to know that the steady state for both mRNA and protein increases by same amount and the nature of the curves remain almost same which is shown in figure 4. In figure 4 the steady state approaches faster than that in figure 3 due to larger degradation rate.

**Figure 4.**
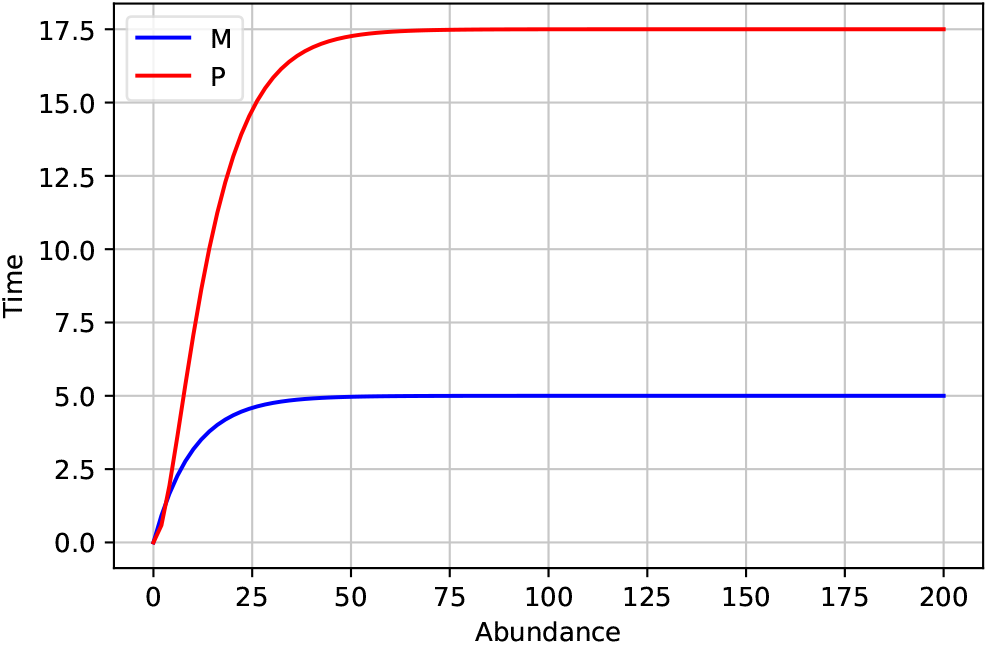
Abundance of mRNA over time with the initial conditions: *k_m_* = 0.5, *γ_m_* = 0.1, *k_p_* = 0.7, *γ_p_* = 0.2

## 3 Hill Function

The so called Hill function were introduced by A.V Hill in 1910 to describe the binding of oxygen to hemoglobin. Subsequently, they have been widely used in biochemistry, psychology and mathematical modeling gene expression^6^. Different mathematical frameworks have been proposed to derive the mathematical model. In particular the use of sets of nonlinear ordinary differential equation(ODE) has been proposed to model the dynamics of the concentrations of mRNAs and proteins. These models are usually characterized by the presence of highly nonlinear Hill function terms^7^. Hill functions follow from the equilibrium state of the reaction in which n ligands simultaneously bind a single receptor^6^.

The Central dogma of molecular biology states that DNA makes and RNA makes proteins. The process by which DNA is copied to RNA is called transcription and by which RNA is used to produce protein is translation. The Hill function is expressed as follow:

- Activation Hill function
- Repression Hill function

### 3.1 Activation Hill Function

In this function Gene first (*G*_1_) acts as an activator for Gene second (*G*_2_) and it increases the probability of transcription often by increasing probability of RNA polymerase binding.

The coupled differential equations representing the model of activation Hill function are shown in equation 3 & equation 4.

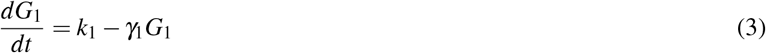

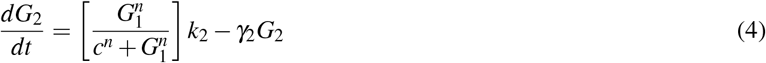

where

- *k*_1_ is production rate of *G*_1_
- *γ*_1_ is degradation rate of *G*_1_
- *k*_2_ is production rate of *G*_2_
- *γ*_2_ is degradation rate of *G*_2_
- c = constant
- n = hill constant

The figures 5 & 6 show the relation between Gene first (*G*_1_) and Gene second (*G*_2_) over time. Along the x-axis, we have the time axis and along y-axis we have number of Gene first (*G*_1_) and Gene second *G*_2_. We observe that *G*_1_ (blue line) denotes Gene first in the above figure 5 & 6 while *G*_2_ (red line) denotes Gene second. Here, (*G*_1_) quickly get activated and reaches to the steady point while *G*_2_ delays in figure 5. This is because for activating the *G*_2_, *G*_1_ should be produced.

**Figure 5.**
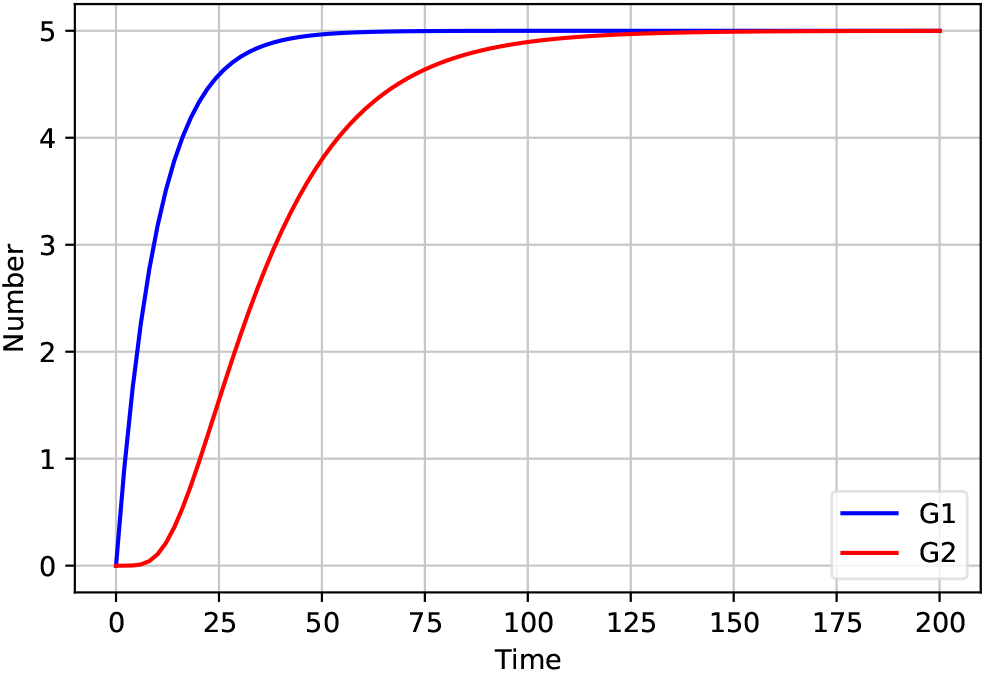
Number of gene over time with the initial conditions: *k*_1_ = 0.5, *γ*_1_ = 0.1, *k*_2_ = 0.5, *γ*_2_ = 0.05, n=5, c=5

If we change the initial parameters such as *k*_1_, *γ*_1_, *k*_2_, *γ*_2_ greater than that are used in figure 5 above we came to know that the steady state for both Gene first (*G*_1_) and Gene second (*G*_2_) changes and the nature of the curves also changes which is shown in figure 6. If *G*_1_ be the zero than the production rate of gene two will also be zero.

**Figure 6.**
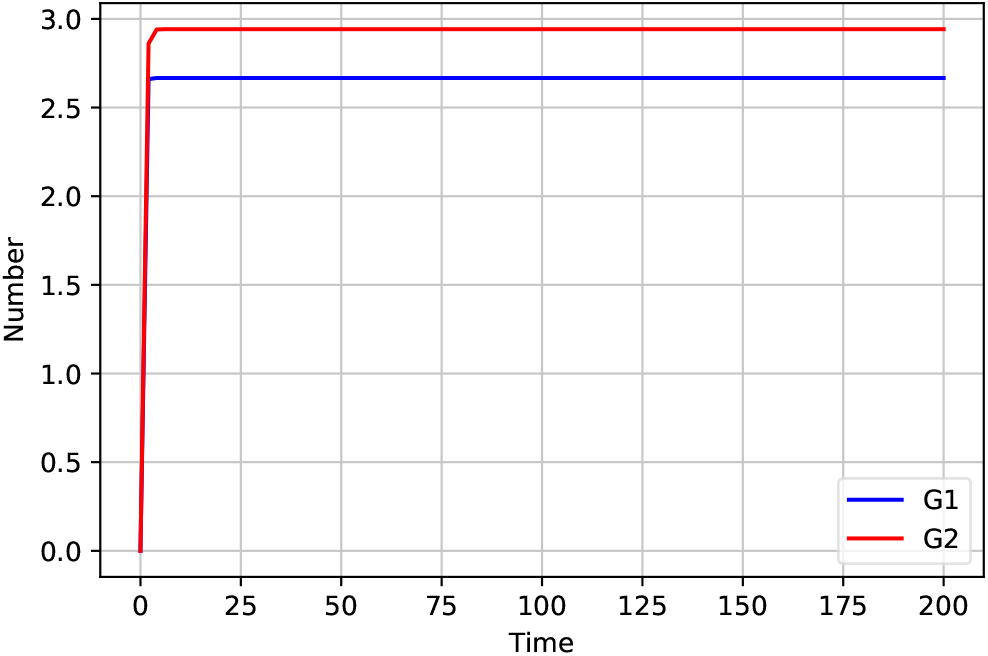
Number of gene over time with the initial conditions: *k*_1_ = 8, *γ*_1_ = 3, *k*_2_ = 6, *γ*_2_ = 2, n=4, c=1

### 3.2 Repression Hill Function

In this function, Gene first (*G*_1_) acts as an repressor for Gene second (*G*_2_) and it decreases the probability of transcription often by decreasing probability of RNA polymerase binding. The damping of protein production by a repressive agent occurs linearly but fluctuations can show a maximum at intermediate repression^8^.

The coupled differential equations representing the model of repression Hill function are shown in equation 5 & equation 6.

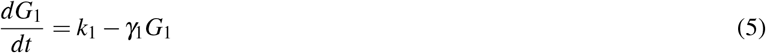

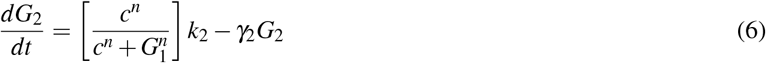

where

- *k*_1_ is production rate of *G*_1_
- *γ*_1_ is degradation rate of *G*_1_
- *k*_2_ is production rate of *G*_2_
- *γ*_2_ is degradation rate of *G*_2_
- c = constant
- n = hill constant

The figures 7 & 8 show the relation between Gene first (*G*_1_) and Gene second (*G*_2_) over time. Along the x-axis, we have the time axis and along y-axis we have number of Gene first (*G*_1_) and Gene second*G*_2_. We observe that *G*_1_(blue line) denotes Gene first in the above figure 7 & 8 while *G*_2_(red line) denotes Gene second. Here ((*G*_2_) quickly get activated and reaches to the peak point but *G*_1_ protein start repressing it and suddenly it goes down in figure 7. This show that *G*_1_ act as repressor in this case.

**Figure 7.**
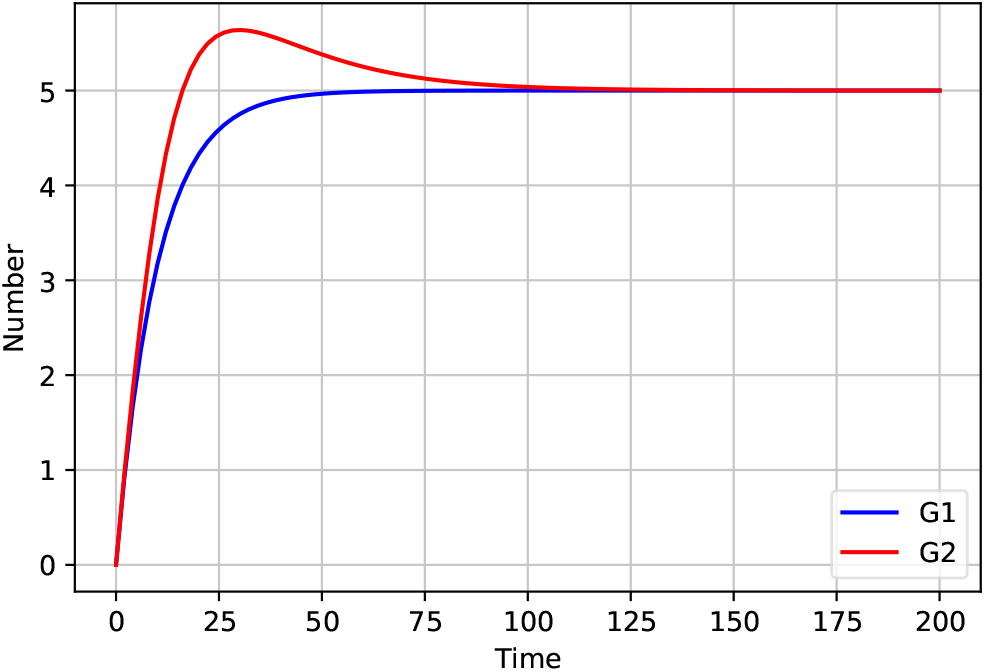
Number of gene over time with the initial conditions: *k*_1_ = 0.5, *γ*_1_ = 0.1, *k*_2_ = 0.5, *γ*_2_ = 0.05, n=5, c=5

**Figure 8.**
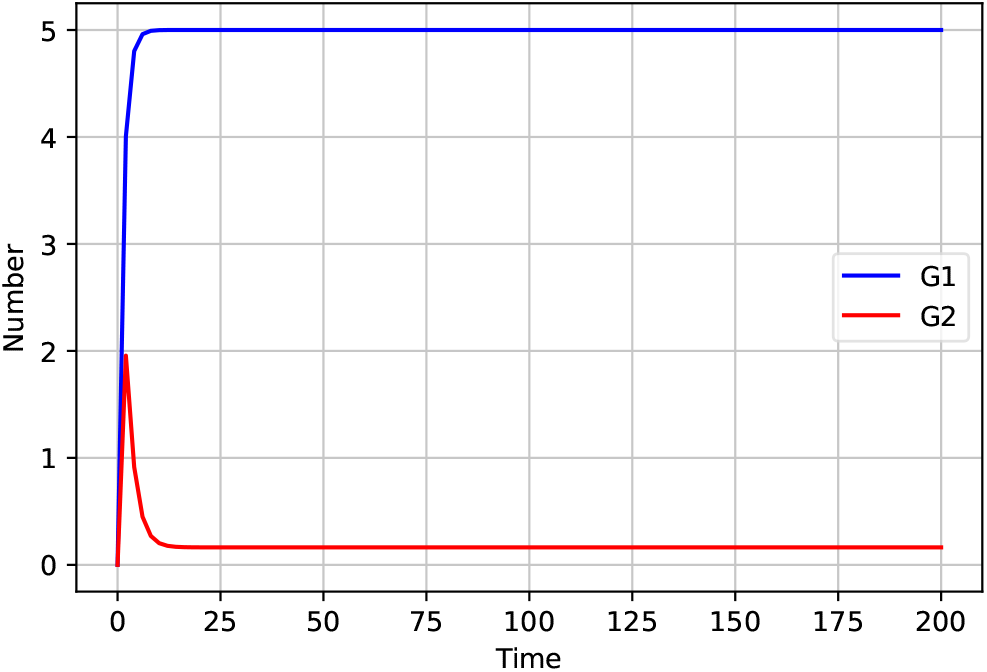
Number of gene over time with the initial conditions: *k*_1_ = 4, *γ*_1_ = 0.8, *k*_2_ = 0.3, *γ*_2_ = 0.5, n=7, c=3

If we change the initial parameters such as *k*_1_, *γ*_1_, *k*_2_, *γ*_2_ greater than that are used in figure 7, we came to know that the Gene first (*G*_1_) approaches near to zero. If *G*_1_ is very large, than the gene two will reach the peak point but *k*_2_ approaches very close to zero but never can be zero.

## 4 Oscillating Gene Network

Oscillating gene is a gene that is expressed in a rhythmic pattern or in periodic cycles^9, 10^. Oscillating genes are usually circadian and can be identified by periodic changes in the state of an organisms. Oscillating gene model is a complex gene network. Same as above, the Central Dogma of molecular biology states that DNA makes and RNA makes proteins. The process by which DNA is copied to RNA is called transcription and by which RNA is used to produce protein is translation.

For this model, let us consider Gene first as (*G*_1_), Gene second as (*G*_2_) and Gene third (*G*_3_). *G*_1_ activates the *G*_2_ and facilitates the transcription of *G*_2_ as a result *G*_2_ get transcribed. It is positive interaction. *G*_2_ does same for the *G*_3_ and facilitates the transcription of *G*_3_ as a result *G*_3_ get transcribed. But *G*_3_ comes back to repress the transcription of *G*_1_. *G*_3_ is inhibiting the expression by blocking the transcription of *G*_1_. It is known as negative feedback because it is cascading *G*_1_ being transcribed and then activating *G*_2_ which later transcribed and helps in activating *G*_3_. But *G*_3_ is negatively feeding back. This cause oscillation during stimulation.

The coupled differential equations representing the model of Oscillating gene network are shown in equation 7, equation 8 & equation 9.

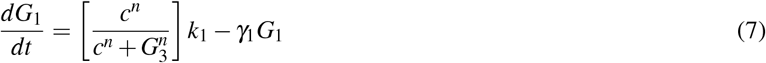

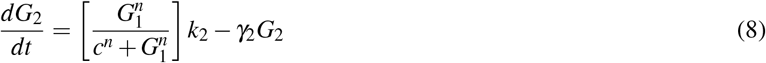

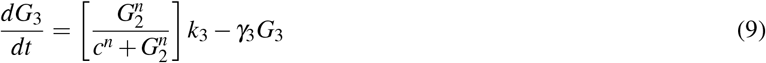

where

- *k*_1_ is production rate of *G*_1_
- *γ*_1_ is degradation rate of *G*_1_
- *k*_2_ is production rate of *G*_2_
- *γ*_2_ is degradation rate of *G*_2_
- *k*_3_ is production rate of *G*_3_
- *γ*_3_ is degradation rate of *G*_3_
- c = constant
- n = hill constant

The figures 9 & 10 show the relation between Gene first (*G*_1_), Gene second (*G*_2_) and Gene third (*G*_3_) over time. Along the x-axis, we have the time axis and along y-axis we have number of Gene first (*G*_1_), Gene second *G*_2_ and Gene third (*G*_3_). We observe that *G*_1_(blue line) denotes Gene first, *G*_2_(red line) denotes Gene second and *G*_3_(green line) denotes Gene third. In figure 9, we find oscillation with 3G network. Gene third *G*_3_ is repressing *G*_1_ which is negative feedback. Here *G*_1_ is being produced that leads to *G*_2_ production and *G*_2_ is produced which leads to the production of *G*_3_. But *G*_3_ produced stops *G*_1_ from being produced. So that we get waves like this.

**Figure 9.**
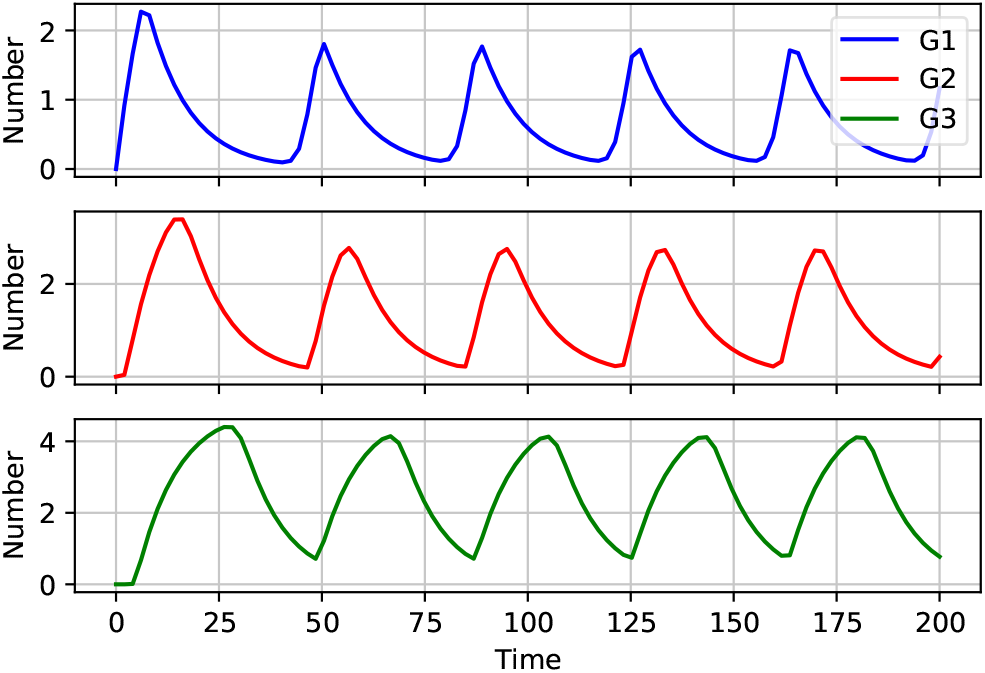
Number of gene over time with the initial conditions: *k*_1_ = 0.5, *γ*_1_ = 0.1, *k*_2_ = 0.5, *γ*_2_ = 0.01, *k*_3_ = 0.5, *γ*_3_ = 0.1, n=9, c=1

If we change the initial parameters such as *k*_1_, *γ*_1_, *k*_2_, *γ*_2_, *k*_3_ and *γ*_3_ greater than that are used in figure 9 came to know that the nature of oscillation of all gene remain same but peak points of the all gene varies from figure 9 for figure 10.

**Figure 10.**
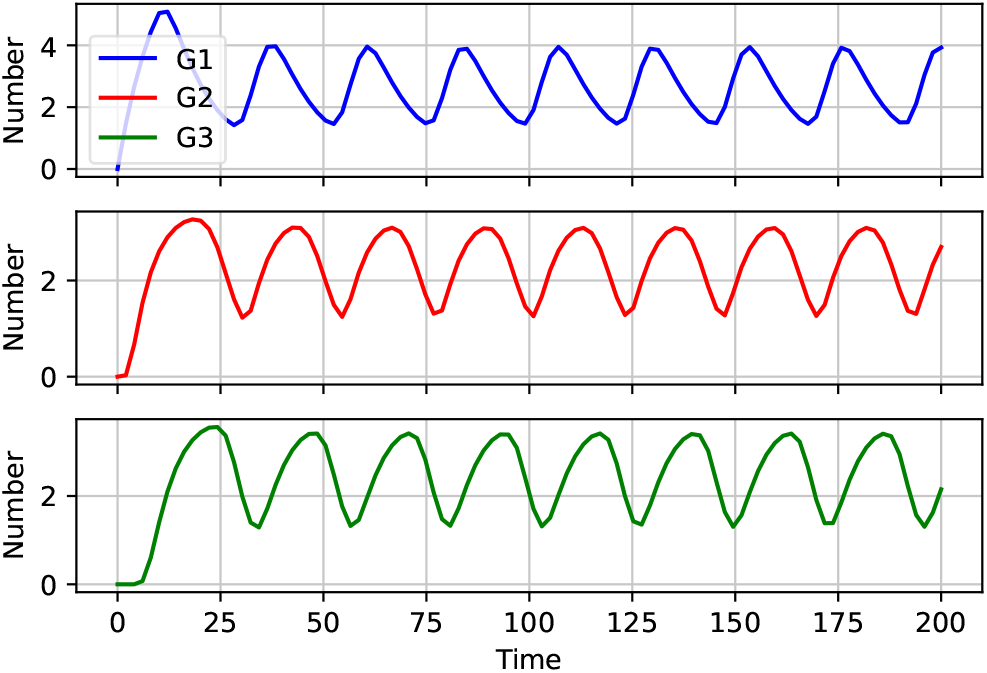
Number of gene over time with the initial conditions: *k*_1_ = 0.8, *γ*_1_ = 0.1, *k*_2_ = 0.7, *γ*_2_ = 0.2, *k*_3_ = 0.8, *γ*_3_ = 0.2, n=6, c=2

## 5 Gillespie Algorithm

The Gillespie algorithm(or occasionally the Doob-Gillespie algorithm) generates a statistically correct trajectory(possible solution) of a stochastic equation system for which the reaction rates are known. The algorithm was first presented by Doob in the mid 1940s^11, 12^. It was implemented by Kendall in the 1950s^13, 14^. However it wasn’t until the mid 1970s^15, 16^ that Gille-spie re-derived the method by studying physical systems that it became widely used^17^. It is also sometime called as SSA method.

For this let consider a gene as x and some mRNA is transcribed from x with production rate k and it will be degraded with rate *γ_x_*. For this list, all the rate in array as r = [k, *γ_x_*]. For this, let choose a time point of the reaction. If we are at time t which could be zero or can be any time between algorithm. Then next time is going to be t + *τ*. Here we choose *τ* from exponential random distribution with the parameter lambda which is the sum of all rates at time t. After choosing the time point it is important to know whether it will be production of mRNa or breakdown of mRNA. The rate for breakdown depends on the current level of x. If there will be x zero than *γ_x_* will be zero and there will be only production. But suppose, if we have couple of x with some positive number. So to choose the event going to happen we take random draw between two of them and weight the probabilities accordingly. Here probability of production of mRNA that means probability of x going to 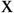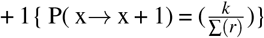 and probability of breaking down of mRNA is that means probability of 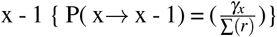.

The differential equation representing the model of repression Gillespie algorithm is shown in equation 10.

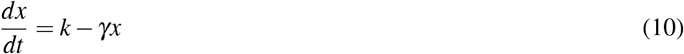

where

- k is the production rate for mRNA
- *γ_x_* is the degradation rate for mRNA

| Events | Rates |
| --- | --- |
| $x \rightarrow x + 1$ | $k$ |
| $x \rightarrow x - 1$ | $\gamma x$ |

The figures 11 & 12 show the relation between quantity of mRNA and protein over time. Along the y-axis, we have the time axis and along x-axis we have abundance of both mRNA and protein. We observe that M(blue line) denotes quantity of mRNA in the above figures 11 & 12. If we do the stimulation for hundred times or thousand times it will show average steady state near about 20 for above figure 11. Since it is stochastic model the steady state is not real because it is fluctuating near the steady state.

**Figure 11.**
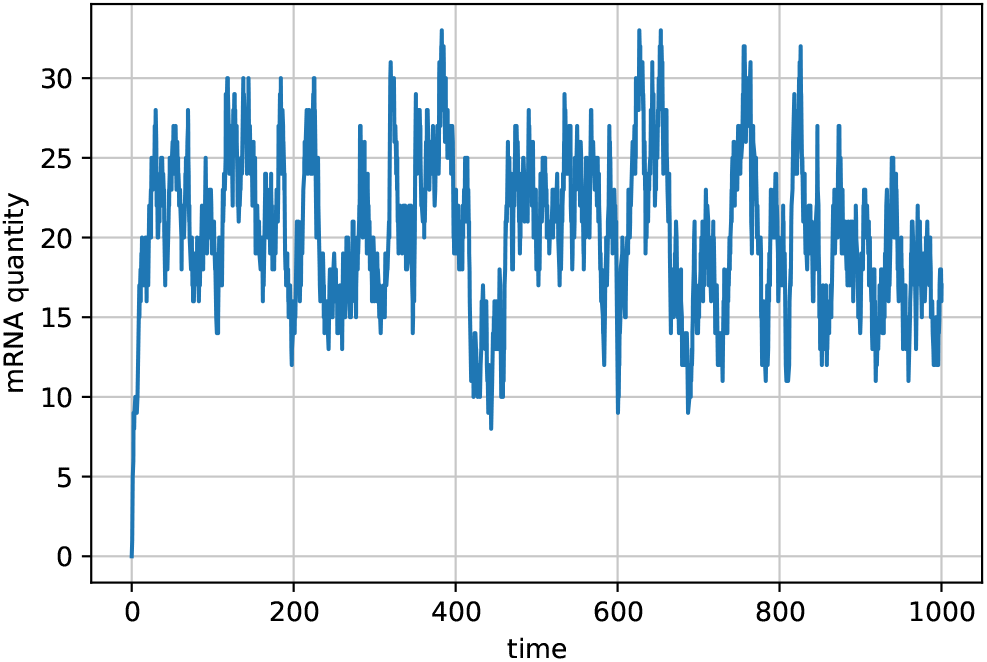
Quantity of mRNA over time with the initial conditions: k=2, *γ* = 0.1

If we change the initial parameters of k and *γ* greater than above figure 11, we came to know that fluctuation is more and hard to find the steady state in figure 12.

**Figure 12.**
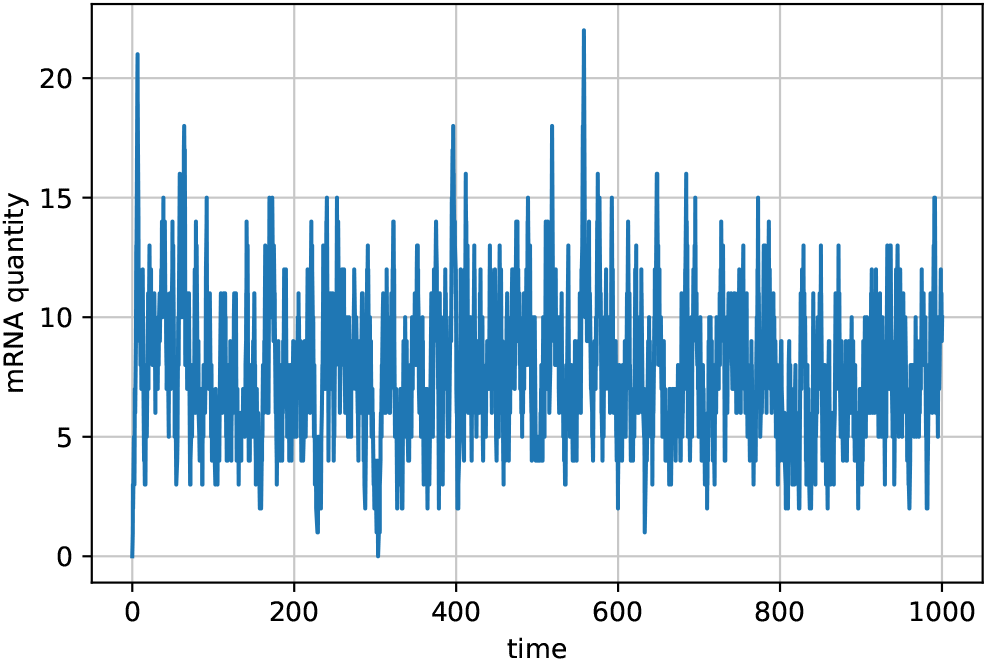
Quantity of mRNA over time with the initial conditions: k=3, *γ* = 0.4

## 6 Deterministic vs Stochastic Modelings

Deterministic modeling allows us to calculate a future event exactly without the involvement of randomness. Stochastic modeling allows us to calculate a future event exactly with the involvement of randomness. In deterministic there is each set of initial condition so there is only one trajectory. Through this model we get smooth curve. The difference with stochastic model is that stochastic model involves some random component. Due to amalgamation of noise from different source, micro-array expression profiles becomes inherently noisy leading to significant impact on the Gene Regulatory Networks (GRN) reconstruction process. Micro-array replicates (both biological and technical), generated to increase the reliability of data obtained under noisy conditions, have limited influence in enhancing the accuracy of reconstruction. Therefore instead of the conventional GRN modeling approaches which are deterministic, stochastic techniques are becoming increasingly necessary^18^.

The trajectory of deterministic modeling is completely defined by the initial parameters and conditions due to no involvement of randomness but in stochastic modeling with the same parameters and same initial conditions we get different trajectories running at different times. In computational biology one motive for using stochastic modeling is to calculate random variation in system which is not possible through deterministic modelings. Here, we will discuss about oscillating gene network same as above for stochastic modeling calculation.

The coupled differential equations representing the model of Stochastic modelings are shown in equation 11, 12 & 13.

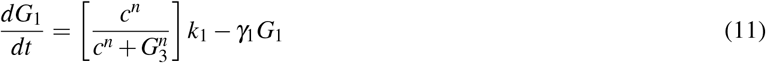

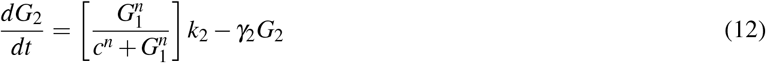

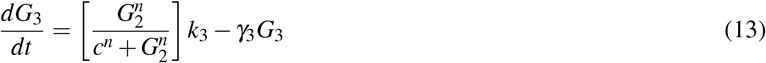

where

- *k*_1_ is production rate of *G*_1_
- *γ*_1_ is degradation rate of *G*_1_
- *k*_2_ is production rate of *G*_2_
- *γ*_2_ is degradation rate of *G*_2_
- *k*_3_ is production rate of *G*_3_
- *γ*_3_ is degradation rate of *G*_3_
- c = constant
- n = hill constant

| Events | Rates |
| --- | --- |
| $G_1 \rightarrow G_1 + 1$ | $\left[ \frac{c^n}{c^n + G_3^n} \right] k_1$ |
| $G_1 \rightarrow G_1 - 1$ | $\gamma_1 G_1$ |
| $G_2 \rightarrow G_2 + 1$ | $\left[ \frac{G_1^n}{c^n + G_1^n} \right] k_2$ |
| $G_2 \rightarrow G_2 - 1$ | $\gamma_2 G_2$ |
| $G_3 \rightarrow G_3 + 1$ | $\left[ \frac{G_2^n}{c^n + G_2^n} \right] k_3$ |
| $G_3 \rightarrow G_3 - 1$ | $\gamma_3 G_3$ |

The figures 13 & 14 show the relation between Gene first (*G*_1_), Gene second (*G*_2_) and Gene third (*G*_3_) over time. Along the x-axis, we have the time axis and along y-axis we have number of Gene first (*G*_1_), Gene second (*G*_2_) and Gene third (*G*_3_). We observe that *G*_1_(blue line) denotes Gene first, *G*_2_(red line) denotes Gene second and *G*_3_(green line) denotes Gene third in both figure 13 & 14. Each one have the oscillations but there is unevenly space because of the stochastic nature of this model. In compared to previous oscillator topic we did deterministic ODEs model which was smooth but this is stochastic so they are still oscillating but it is much more random.

**Figure 13.**
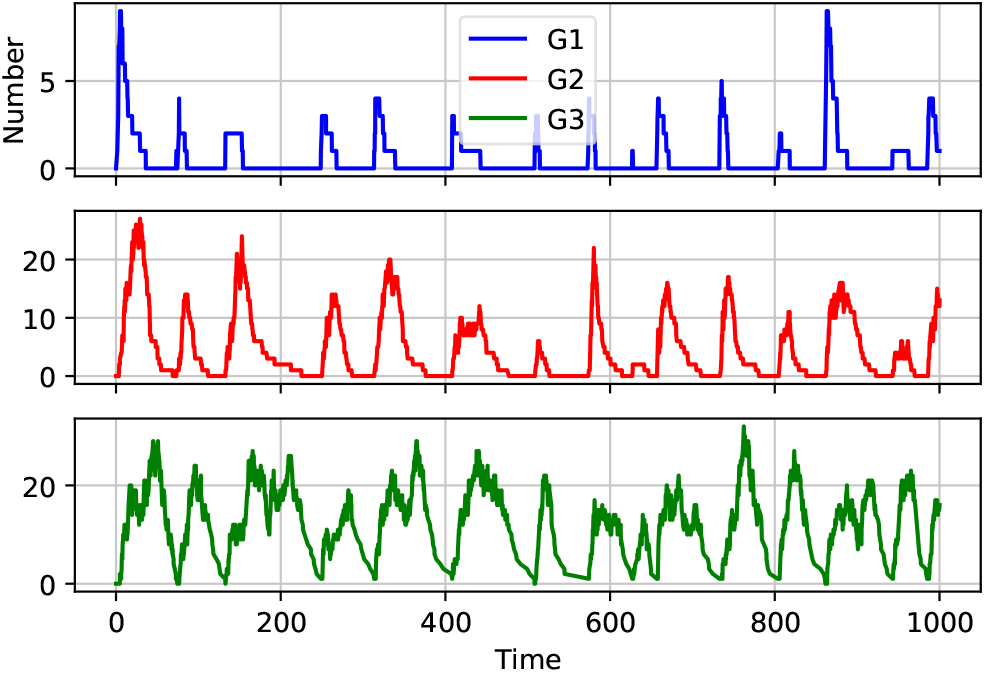
Number of gene over time with the initial conditions: *k*_1_ = 2, *γ*_1_ = 0.1, *k*_2_ = 2, *γ*_2_ = 0.1, *k*_3_ = 2, *γ*_3_ = 0.1, n=9, c=1

If we change the initial parameters such as *k*_1_, *γ*_1_, *k*_2_, *γ*_2_, *k*_3_ and *γ*_3_ greater than the above figure 13, we came to know that the nature of oscillation of all Gene remain same but peak points of the all Gene varies from figure 13 and some what oscillates in rapid speed in less time in figure 14.

**Figure 14.**
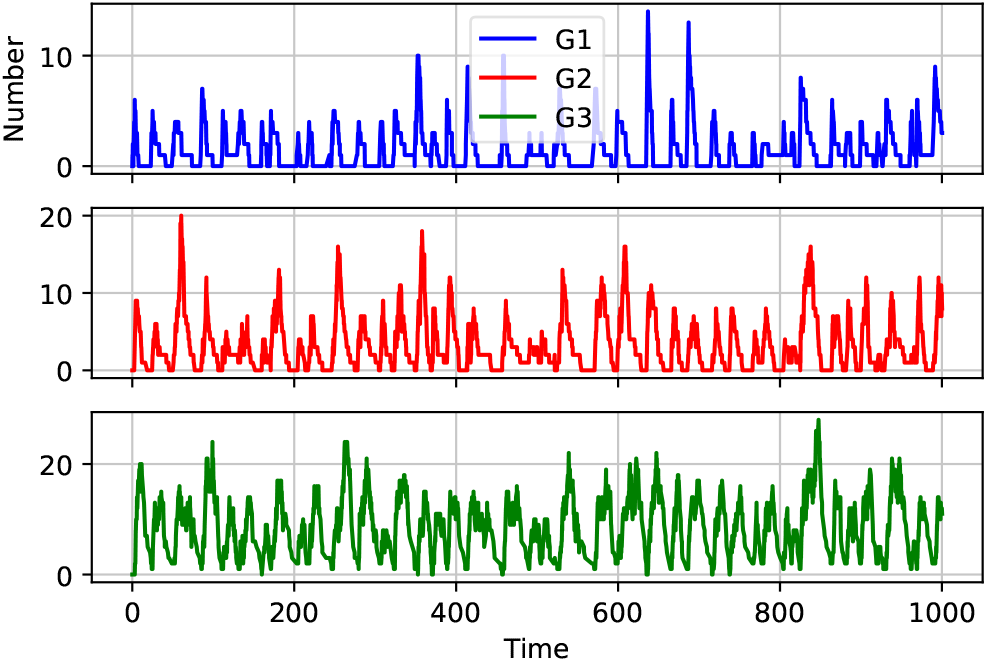
Number of gene over time with the initial conditions: *k*_1_ = 3, *γ*_1_ = 0.2, *k*_2_ = 2.5, *γ*_2_ = 0.2, *k*_3_ = 3, *γ*_3_ = 0.1, n=10, c=2

## 7 Discussion

As mentioned above, this manuscript has been engaged in the use of mathematical modelling in the Gene Regulatory System (GRN). Here, we underlined that the model should describes the continuous nature of the biochemical processes and reflect the non-linearities. We used the gene expression rates by ordinary differential equations (ODEs) where the expression rates were approximated by means of difference quotients. For the modeling, Python modules were used. Python has many excellent and well-maintained libraries that facilitate high-level scientific computing analyses. The modules such as numpy^19^ which offers a numerical processing library that supports multi-dimensional arrays, scipy^20^ which offers a scientific processing library providing advanced tools for data analysis, including regression, ODE solvers and integrator, linear algebra and statistical functions, matplotlib^21^ which offers a plotting library with tools to display data in a variety of ways were used in Anaconda^22^ software. The dynamical behavior of gene expressions which are represented by a system of ordinary differential equations were solved using “odeint” from scipy module. Further we also considered the different parameters to solve the different ODEs. Finally, the change of dependent variable with respect to time were plotted in different models under study.

In Central dogma, we observe that flow of genetic information in a biological system is steady after a certain time, and the abundance of both mRNA and protein depends on the production. Likewise in Hill function we observe that it is a parameters which increases or decreases the probability of RNA polymerise binding. In Oscillating gene network, we observe that it is a positive complex network also known as negative feed back. In Gillespie algorithm, we observe that it is an algorithm for predicting the time point of the reaction between the mRNA and proteins. In the Deterministic vs Stochastic modelings, we observe that deterministic model can calculate the future event with out randomness but stochastic can calculate future with randomness. We also discussed about the randomness of gene over a time which causes the significant impact on the Gene Regulatory Network (GRN). The Stochastic model is far better than Deterministic model to calculate future event exactly with low chance of error. We are able to predict the future event using different parameters. Thus, modeling is a mathematical tool, like a microscope, which allows consequences to logically follow from a set of assumptions. It may allow to understand importance of specific mechanisms/assumptions in biological processes. However, mathematical modeling can also be misleading if used inappropriately.

## Appendix

The code that was used in order to study different types of modeling in biological systems are shown here. This will help to reproduce the solutions to the models discussed above.

## Central Dogma

The code for the Central Dogma problem that is used in order to generate the plots is shown here:

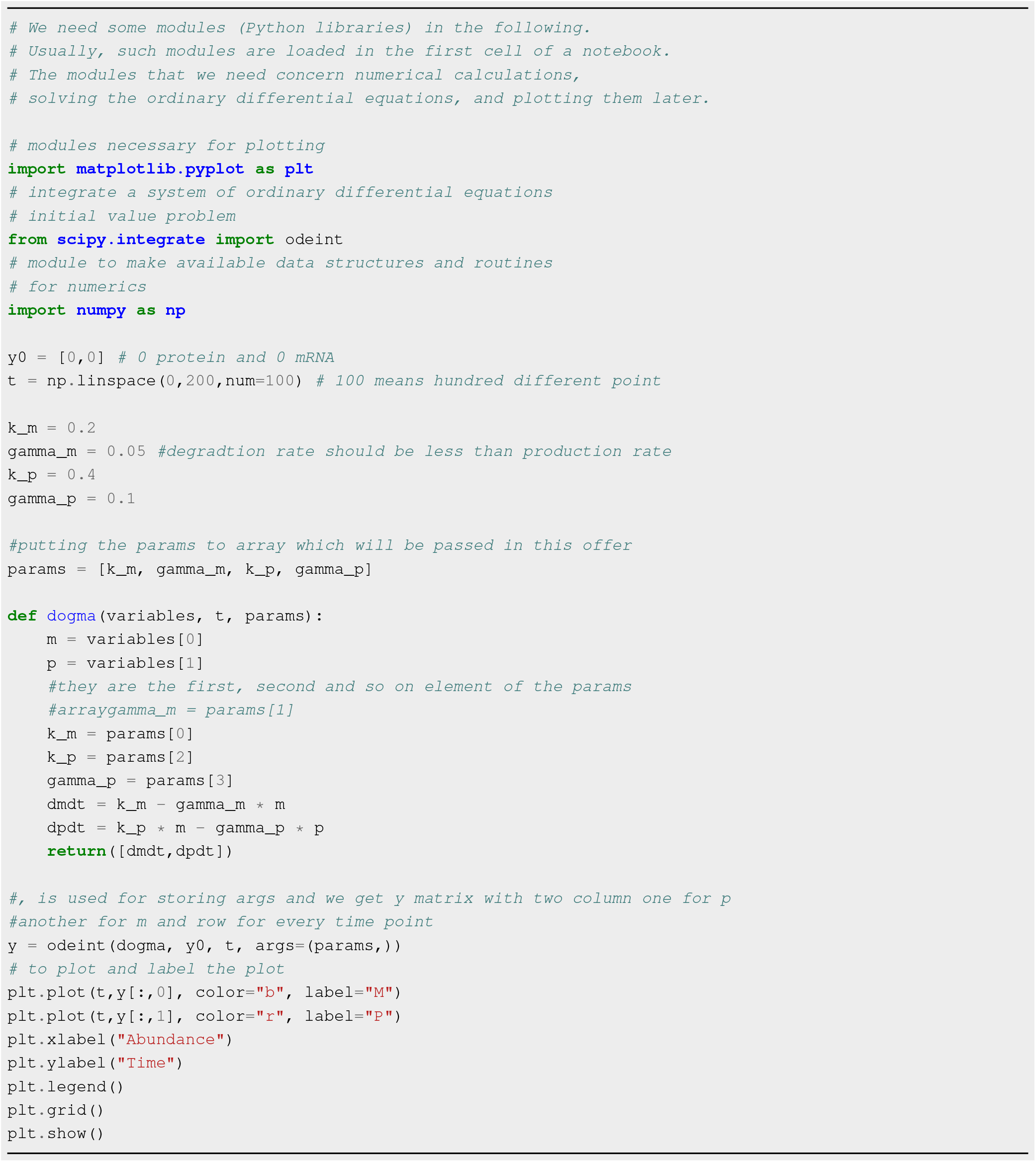

## Hill function

The code for the Hill function problem that is used in order to generate the plots is shown here:

## Activation Hill Function

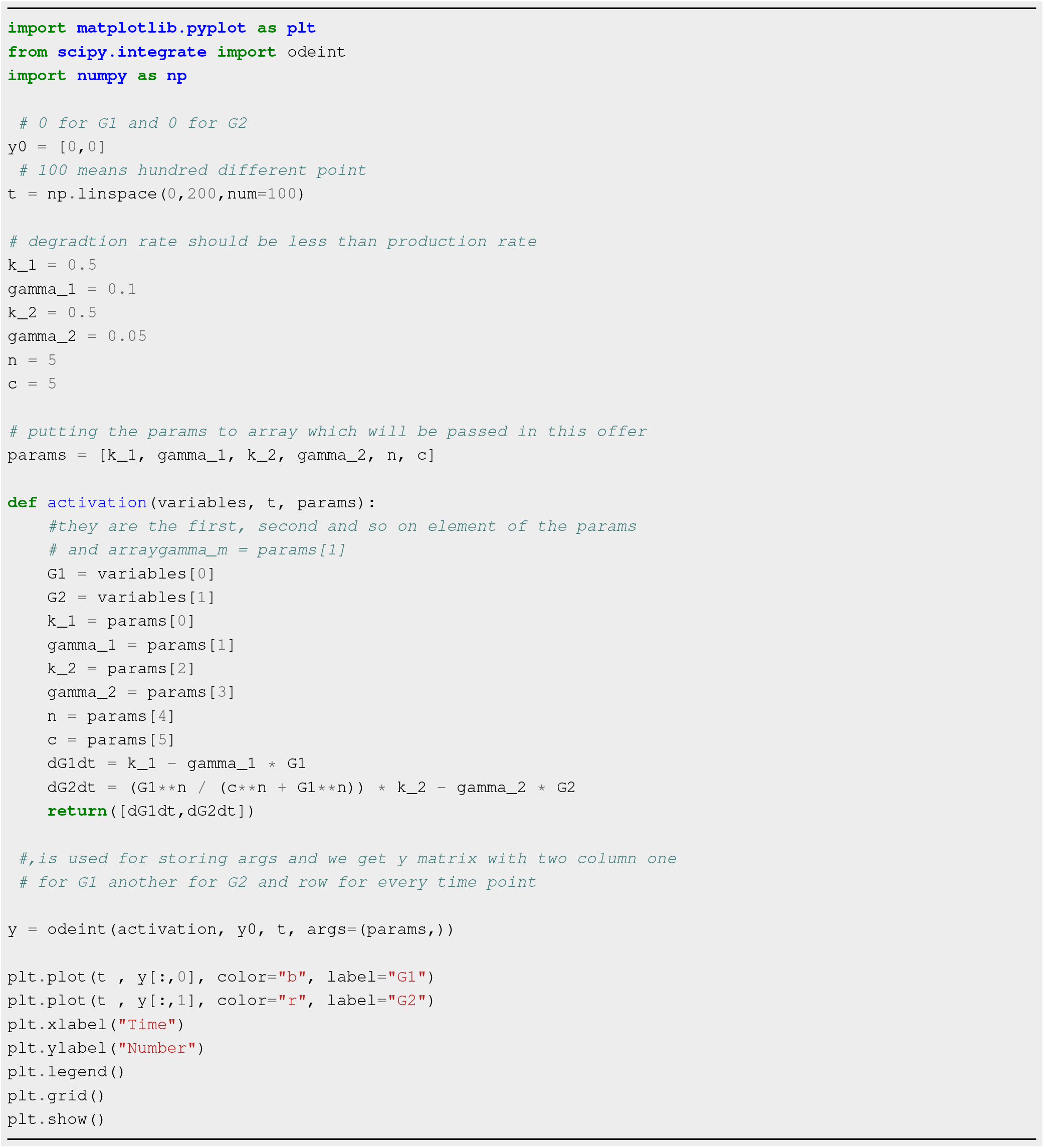

## Repression Hill Function

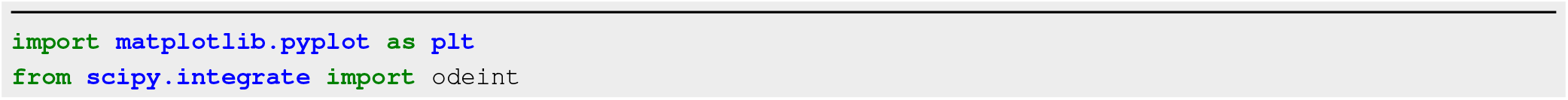
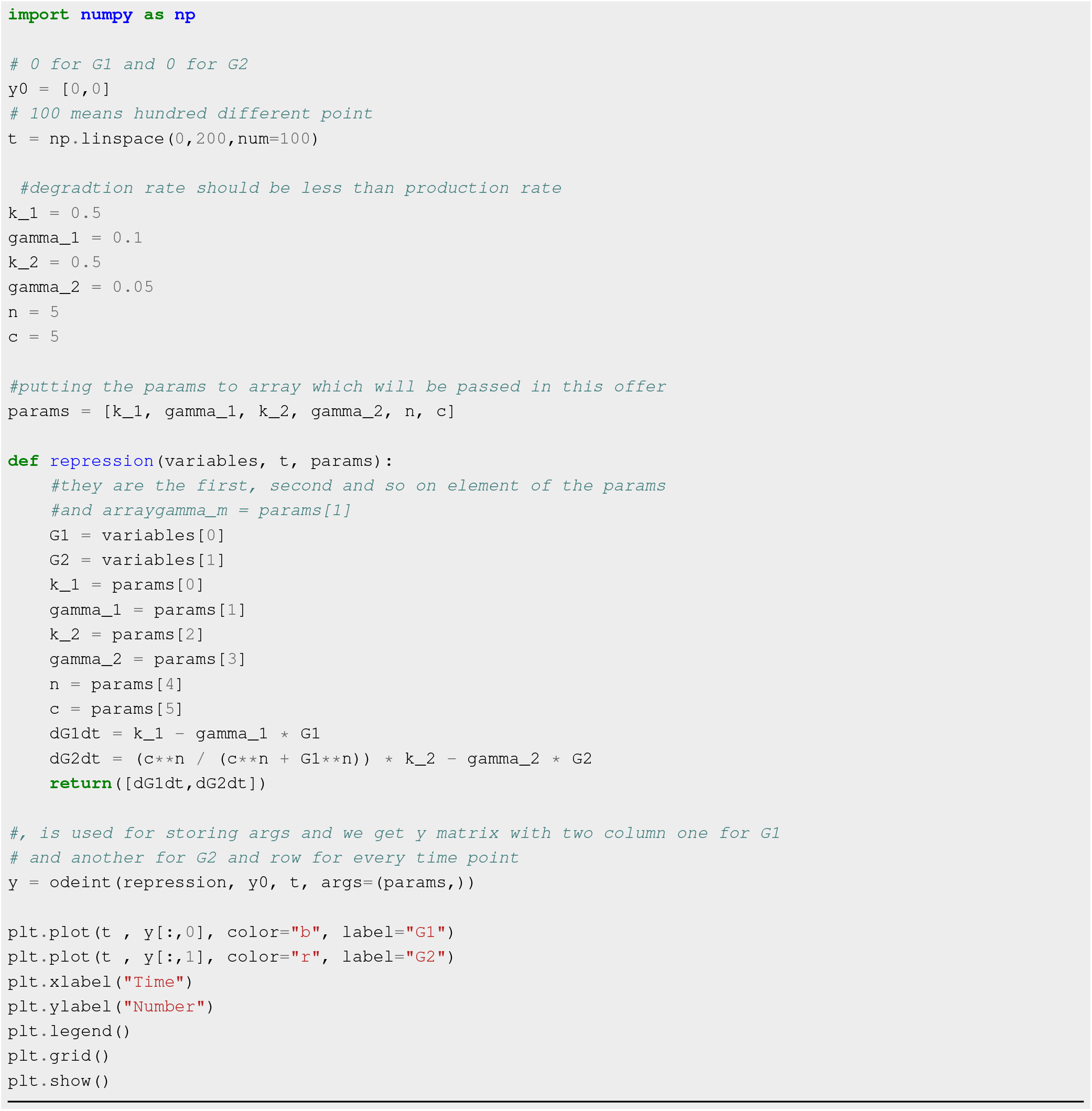

## Oscillating Gene Network

The code for the Oscillating gene network problem that is used in order to generate the plots is shown here:

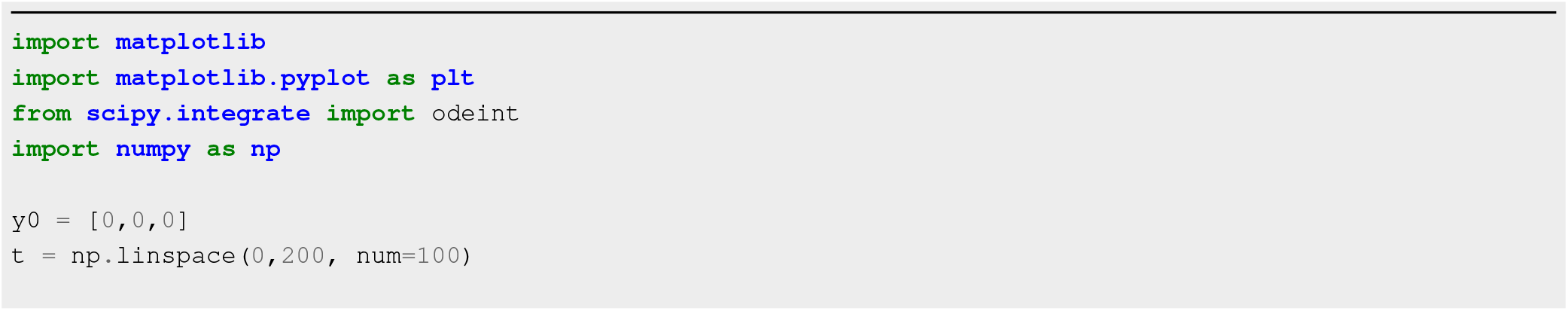
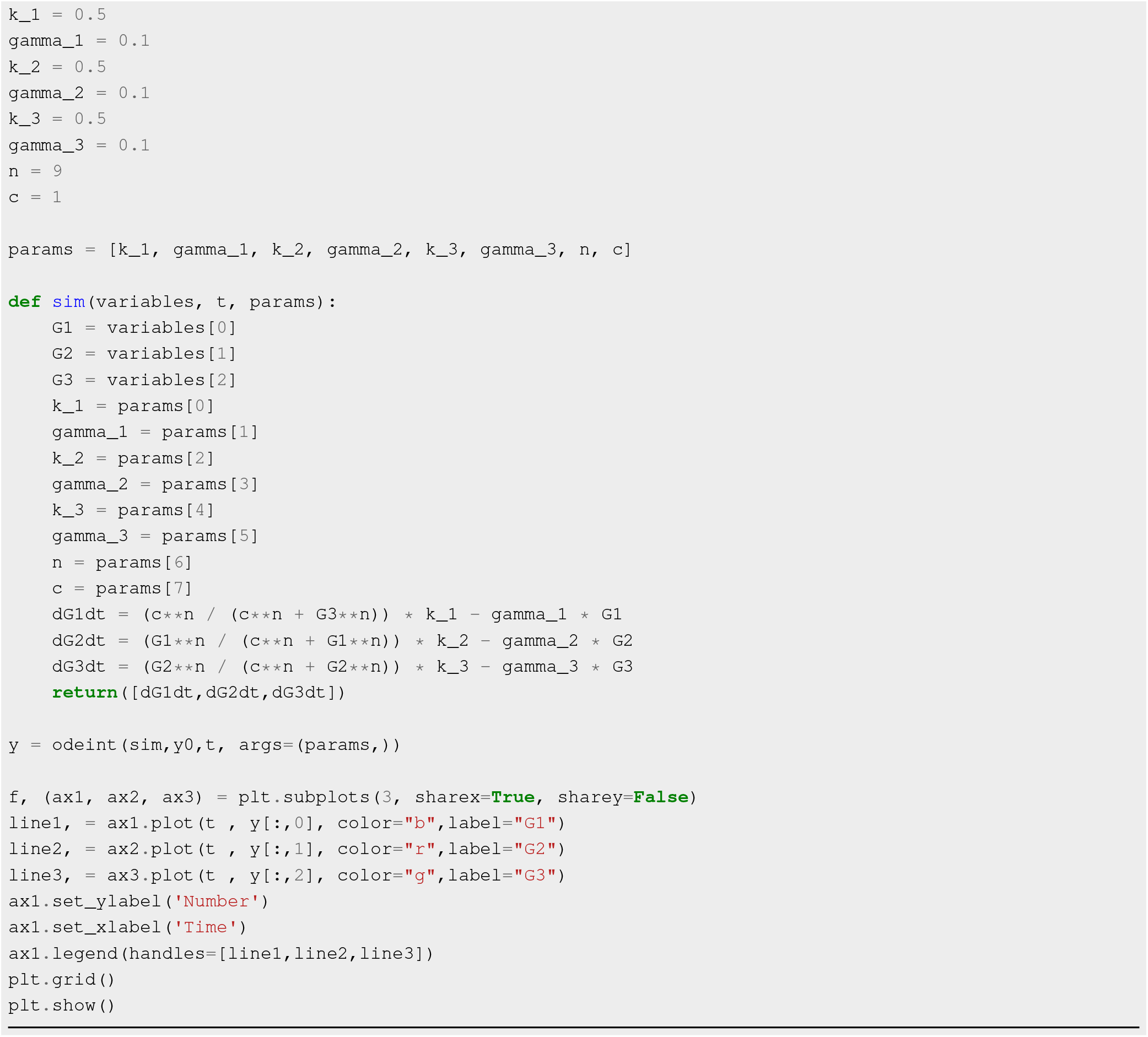

## Gillespie Algorithm

The code for the Gillespie algorithm that is used in order to generate the plots is shown here:

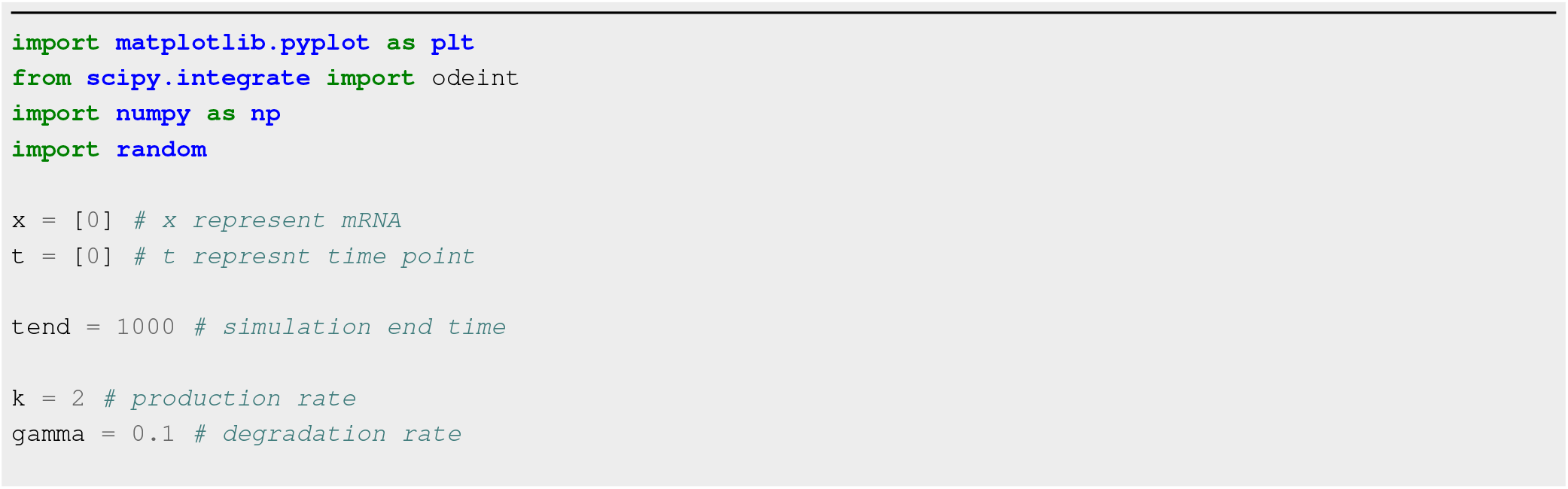
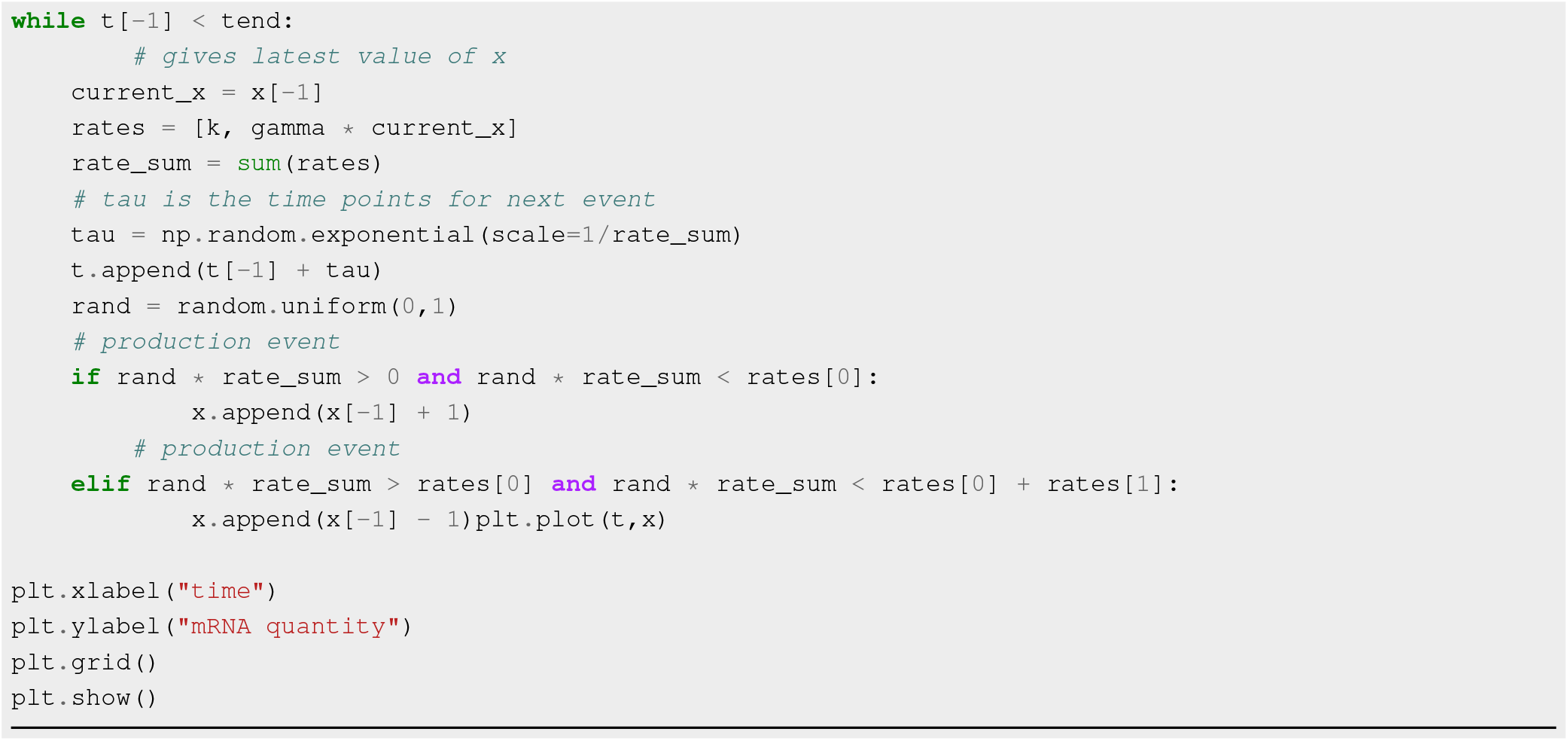

## Deterministic vs Stochastic modelings

The code for the Deterministic vs Stochastic modelings problem that is used in order to generate the plots is shown here:

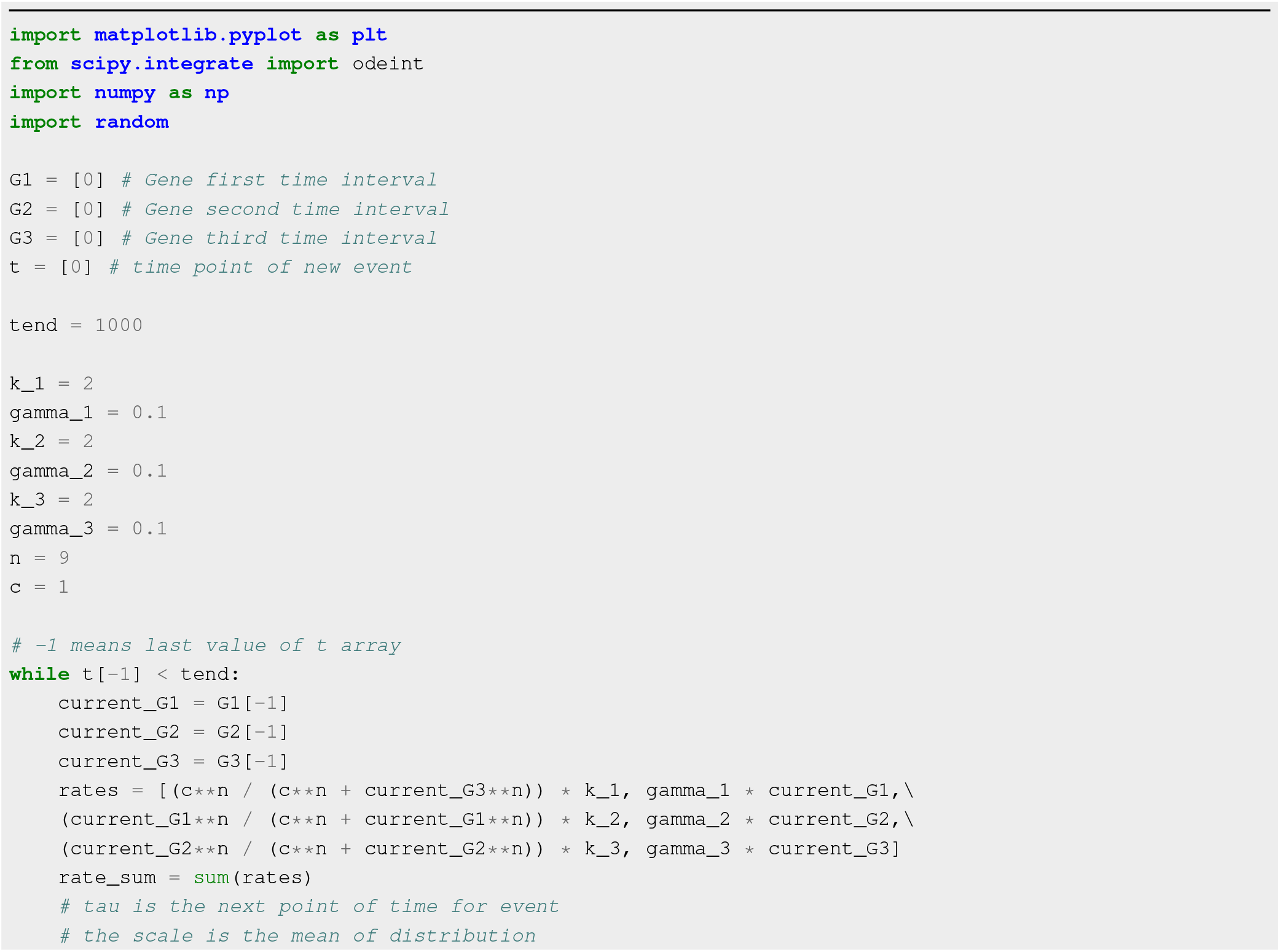
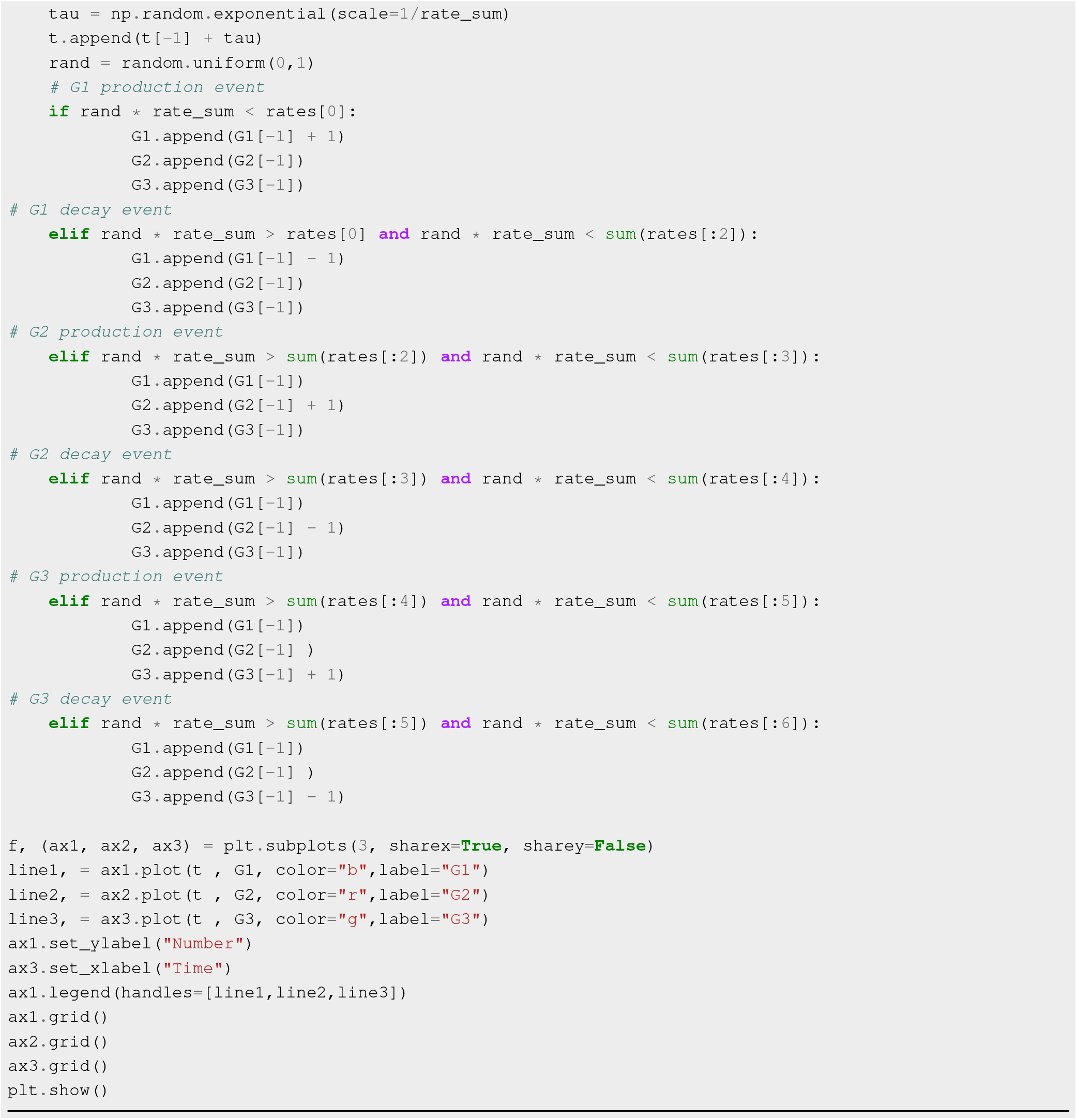

## Footnotes

1 github.com/aaditya-pdgupta/mathematical-modeling-in-biology

